# Fe(III)-dependent anaerobic methane-oxidizing activity in a deep underground borehole demonstrated by in-situ pressure groundwater incubation

**DOI:** 10.1101/2022.01.27.478104

**Authors:** Hiroki Nishimura, Mariko Kouduka, Akari Fukuda, Toyoho Ishimura, Yuki Amano, Hikari Beppu, Kazuya Miyakawa, Yohey Suzuki

## Abstract

The family Methanoperedenaceae archaea mediate anaerobic oxidation of methane (AOM) in various terrestrial environments. In this study, we newly developed a high-pressure laboratory incubation system by controlling hydraulic pressure ranging from ambient to 5 MPa. Using the system, we investigated groundwater from 214- and 249-m deep boreholes at Horonobe Underground Research Laboratory, Japan, where the high and low abundances of Methanoperedenaceae archaea have been revealed by genome-resolved metagenomics, respectively. We incubated the groundwater samples amended with or without amorphous Fe(III) as an electron acceptor and ^13^C-labelled methane at an in-situ pressure of 1.6 MPa. After three to seven-day incubation, AOM activities were not detected from the 249-m deep groundwater but from the 214-m deep groundwater. The AOM rates were 93.7 ± 40.6 and 27.7 ± 37.5 nM/day with and without Fe(III) amendment. To clarify the differences in AOM activity between the 214- and 249-m deep groundwater samples, we characterized Fe(III) contents in suspended particulates collected by filtration. The particulates were not visible in the 249-m deep groundwater on the filter, while they were abundant and contained Fe(III)-bearing phyllosilicates in the 214-m deep groundwater. These results support the in-situ activity of Fe(III)-dependent AOM in the deep subsurface borehole.

## Introduction

Anaerobic oxidation of methane (AOM), a metabolism conducted by microorganisms including anaerobic methanotrophic archaea (ANME) inhabiting deep seafloor sediments, has been long investigated (e.g., Hinrichs *et al*., 1999; Boetius *et al*., 2000). The process of AOM is theoretically predicted to be pressure-dependent, given the increase in dissolved methane concentration with pressure (Timmers *et al*., 2015). Experimentally, the positive correlation between incubation pressure and AOM rate from deep-sea sediments enriched with ANME has been demonstrated by pressure-variable incubation systems (Nauhaus *et al*., 2002; Krüger *et al*., 2008). In the terrestrial subsurface, metabarcoding and metagenomic analyses have revealed the dominance of the family *Candidatus* (*Ca*.) Methanoperedenaceae (formerly ANME-2d; e.g., Miettinen *et al*., 2018; Flynn *et al*., 2013). Some members of *Ca*. Methanoperedenaceae are dominantly found in near-surface environments using versatile electron acceptors such as nitrate (Haroon *et al*., 2013), Fe(III) (Cai *et al*., 2018) and Mn(IV) (Leu *et al*., 2020b). In contrast to the near-surface members demonstrated for their AOM activities by incubation experiments under ambient pressure, subsurface members of *Ca*. Methanoperedenaceae have been poorly characterized for their AOM activities.

Horonobe Underground Research Laboratory (URL) is constructed in the northern Hokkaido, Japan. Our previous metagenomic study of groundwater from the 214-m deep underground borehole has revealed the abundance of *Ca*. Methanoperedenaceae archaea (Hernsdorf *et al*., 2017). The near-complete genome of *Ca*. Methanoperedenaceae reconstructed from the 214-m deep groundwater was equipped with a set of genes required for AOM and potentially involved in reduction of Fe(III) and Mn(IV) (Ettwig *et al*., 2016; Kletzin *et al*., 2015). In contrast, the abundance was not detected in the 249-m subsurface of the URL where the groundwater has similar hydrogeochemical characteristics with those of the 214-m deep groundwater. Thus, the factors controlling the abundance and activity of *Ca*. Methanoperedenaceae archaea remain largely unknown.

In this study, firstly we conducted 16S rRNA gene amplicon analysis using the 214-m deep groundwater to confirm the dominance of the *Ca*. Methanoperedenaceae archaea reported from our previous study (Hernsdorf *et al*., 2017). Then, we measured AOM rates under in-situ high-pressure conditions by extending previously established incubation experiments with ^13^C-labelled methane (Ino *et al*., 2018). As a result, we succeeded in measuring AOM rates with or without amendments of amorphous Fe(III) in the 214-deep groundwater enriched with the *Ca*. Methanoperedenaceae archaea.

## Results

### Study site and sampling

The Horonobe area is located on the eastern margin of a Neogene to Quaternary sedimentary formations. In September 2019 and September 2021, groundwater samples were collected from an interval of the borehole denoted as 08-E140-C01 which was drilled diagonally downward with vertical angle of −45 degree from the 140 m gallery at Horonobe URL (45°02’43”N, 141°51’34”E). The average interval depth was 214 m below ground level (m b.g.l.). Hereafter, this groundwater is referred to as ESB214. In September 2021, the other groundwater from 249 m b.g.l. (hereafter, referred to as VSB249) was also obtained from the borehole named 09-V250-M02 drilled horizontally from the 250 m gallery. These two boreholes drilled in Koetoi Formation (Neogene siliceous mudstones) were associated with physico-chemically similar groundwater (Table 1). The temperature and hydrostatic pressure are 18 – 20 °C and 1.6 MPa. The geochemical characteristics of groundwater are characterized by neutral pH, salinity lower than seawater and relatively high concentrations of total organic carbon (TOC) around 20 – 50 mg/L (Miyakawa *et al*., 2020). The groundwater from both depths are nearly depleted in electron acceptors for AOM such as nitrate and Fe(III) and contain methane as a main dissolved gas component (90-100%; Miyakawa *et al*., 2017).

**Table 1.**
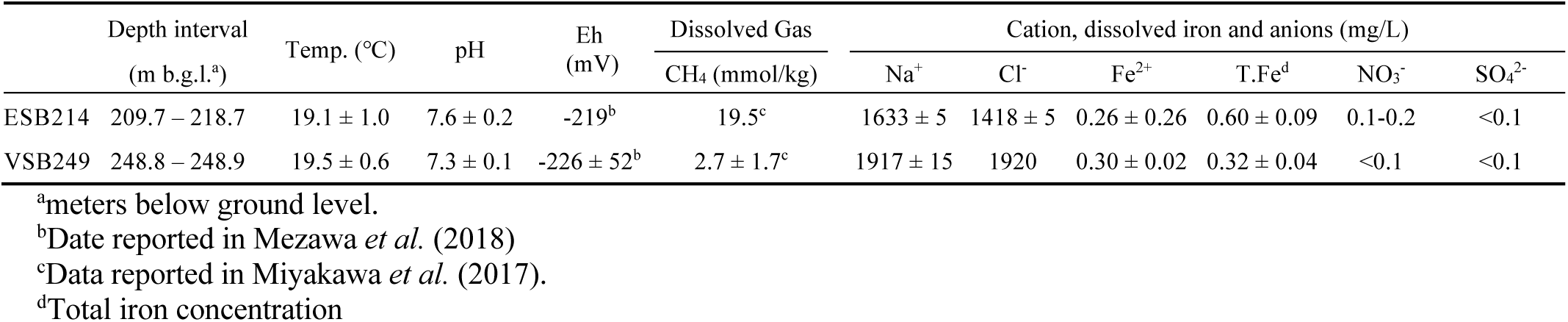
Borehole and groundwater characteristics of Horonobe URL. ESB214 and VSB249 are named for groundwater samples from 08-E140-C01 and 09-V250-M02. All data except for dissolved gas are reported in Miyakawa *et al*. (2020).

To collect groundwater from the target intervals, multi-packer systems were used (Nanjo *et al*., 2012). A pressure-resistant stainless filter holder with a membrane filter was directly connected to the tubing outlet to collect microbial cells without lowering hydrostatic pressure. In-situ hydraulic pressure was retained inside the filter holder during the refrigerated shipment to the laboratory.

### Microbial composition in ESB214

16S rRNA gene amplicon analysis was conducted for ESB214 which was collected in September 2019. We revealed the dominance of the family *Ca*. Methanoperedenaceae, accounting for 10.8% of the total high-quality reads (Fig. 1A). This result confirmed the dominance of *Ca*. Methanoperedenaceae in ESB214 also reported by Hernsdorf *et al*. (2017). The most abundant 16S rRNA gene sequence was affiliated with the genus *Ferribacterium*, the members of which are Fe(III) reducers and obligate anaerobes (Cummings *et al*., 1999). No sequences were related to *Ca. Methylomirabilis oxyfera* (*Ca*. division NC10), the other microbe than *Ca*. Methanoperedenaceae known for AOM (nitrite dependent) at terrestrial settings (Ettwig *et al*., 2010).

**Fig. 1.**
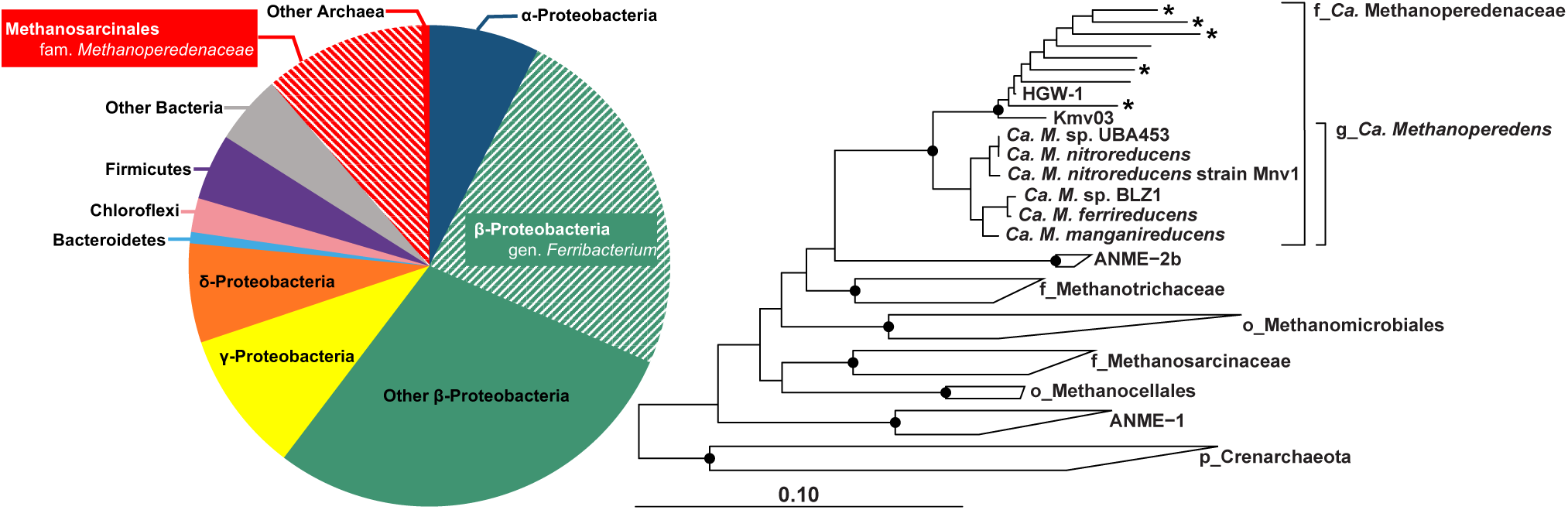
16S rRNA gene sequence analysis of ESB214 collected in 2019. Taxonomic distribution of ESB214 (left). Each color represents major taxonomic groups ranging from genus to phylum level. Taxonomic groups with high proportions are highlighted with shading. Neighbor-joining phylogenetic tree mainly including the family *Ca*. Methanoperedenaceae with sequences obtained from E214 groundwater and public databases (right). This tree was rooted to *Escherichia coli* (not shown). Sequences obtained from ESB214 are indicated without any label (<50 reads) and with asterisks (≥50 reads). The 1000 bootstrap replicates were obtained with a maximum likelihood method and values >80% are shown by gray circles at branches. The scale bar indicates 0.10 expected change per site.

The phylogenetic relationships of *Ca*. Methanoperedenaceae-affiliated sequences were characterized in a neighbor-joining tree (Fig. 1B). The obtained sequences are shown to be diverse within the family *Ca*. Methanoperedenaceae and closely related to HGW-1 and Kmv03, metagenome-assembled genomes (MAGs) reconstructed from ESB214 (Hernsdorf *et al*., 2017) and an active mud volcano (Mardanov *et al*., 2020), respectively.

### Metabolic estimation from closely related Ca. Methanoperedenaceae MAGs

A complete gene set for reverse methanogenesis and related cofactor enzymes are encoded in both HGW-1 and Kmv03 (Leu *et al*., 2020a; Mardanov *et al*., 2020). In the family *Ca. Methanoperedenaceae*, the wide range of electron acceptors is reported for AOM including nitrate (Haroon *et al*., 2013), nitrite (Arshad *et al*., 2015), Fe(III) (Ettwig *et al*., 2016; Cai *et al*., 2018), Mn(IV) (Ettwig *et al*., 2016; Leu *et al*., 2020b), Cr(VI) (Lu *et al*., 2016). HGW-1 and Kmv03 are not equipped with genes involved in the reduction of nitrate but heme-rich multiheme *c*-type cytochromes (MHCs) for extracellular electron transfer. Remarkably, HGW-1 and Kmv03 are equipped with 11 and 18 copies of MHCs with >50 and 22 for heme-binding motifs (CXXCH) in a single MHC gene, respectively. Numerous MHCs with high numbers of heme-binding motifs are commonly found in bacterial groups such as the family Shewanellaceae and Geobacteraceae for their grow by transferring electrons to solid metal oxides (Methé *et al*., 2003; Shi *et al*., 2007; Shi *et al*., 2016). The *Ca. Methanoperedenaceae* members represented by 16S rRNA gene sequences obtained in this study are estimated to perform electron transfers to solid metal oxides in the deep borehole.

### Development of a high-pressure incubation system

For the rate measurements of AOM, a high-pressure incubation system was newly developed to reproduce in-situ borehole conditions. The system was composed of a stainless vessel, a piston, a pressure gauge, valves and gas lines (Fig. 2). Additionally, the medium was incubated in Tedlar bags to avoid direct contact with the vessel material.

**Fig. 2.**
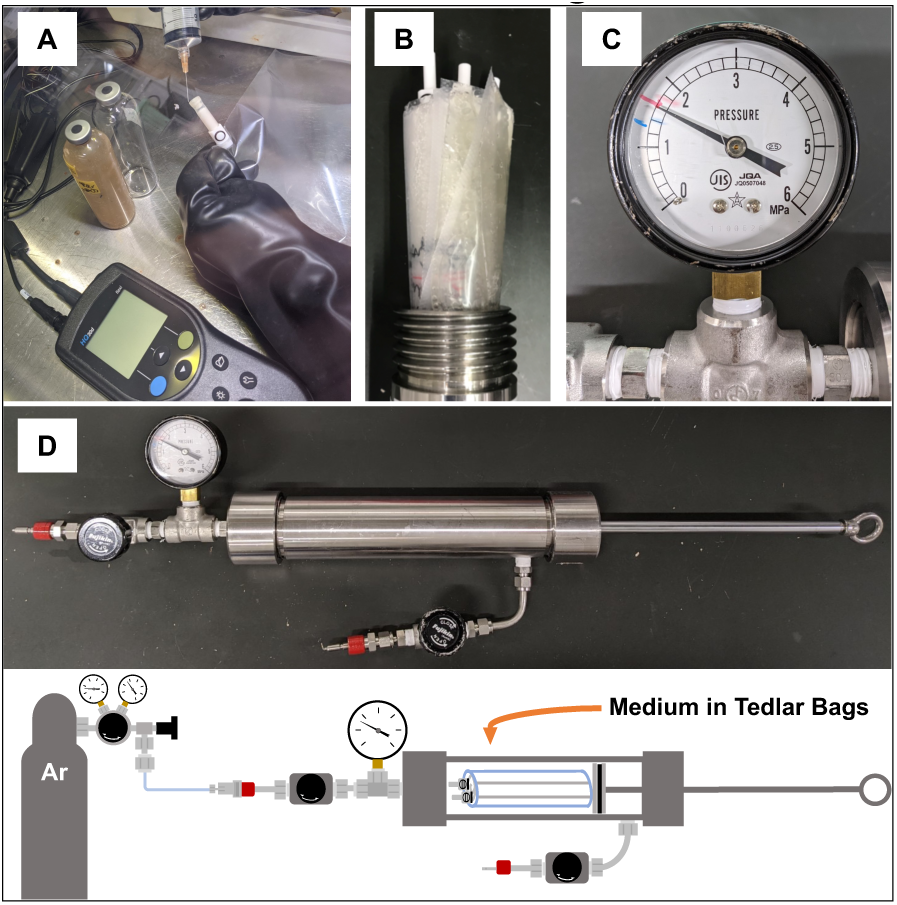
Graphical descriptions of high-pressure incubation experiments. A picture showing medium preparation with a Tedlar bag inside a glovebox (A). The insertion of Tedlar bags into a stainless vessel (B) and a pressure gauge measuring an internal vessel pressure of 1.6 MPa (C). A picture and schematic illustration of the high-pressure incubation system showing the inner vessel structure and the overall experimental setup (D).

Stainless holders, in which in-situ hydraulic pressure was retained for one week after on-site sampling, were opened in a glovebox filled with Ar gas. Microbial cells trapped on the membrane filter inside the stainless holders were suspended in autoclaved, deoxygenated 50 mM NaCl solution (pH 7.4). This NaCl concentration was selected to mimic the salinity of groundwater (Table 1). The original groundwater was not used as a base medium, because the high concentration of dissolved inorganic carbon (DIC) lowers the sensitivity of quantification of ^13^CO_2_ production. Alternatively, a medium was prepared by the ten-fold dilution of filter-sterilized groundwater samples with the 50 mM NaCl solution. Then, the cell suspension was amended either with 0.1 mM amorphous Fe(III), 0.5 mM Na nitrate or 0.5 mM Na sulfate for an artificial electron acceptor. The low levels of O_2_ in the cell suspensions were assured by adding stock FeCl_2_ solution for a final concentration of 2.5 mg/L (scavenge trace amounts of O_2_) and then monitored by a fiber-optic oxygen meter. Finally, the cell suspensions were amended with ^13^C-labeled methane (9 mM) at a concentration nearly equivalent to the groundwater samples (Table 1). Then, the bags were transferred into the stainless vessel and pressurized with Ar gas at an in-situ hydraulic pressure of 1.6 MPa (Fig. 2B, C and D). After three or seven days, the cell suspensions were collected and subjected to chemical and microbiological characterizations.

### AOM activities in borehole groundwater samples

We measured the AOM rates of microbial communities in ESB214 and VSB249 (Fig. 3), where the abundances of *Ca*. Methanoperedenaceae archaea are significantly different (Hernsdorf *et al*. 2017). The changes in carbon stable isotopic ratio (*δ*^13^C) in DIC and alkalinity during incubation were measured to calculate the AOM rates. Microbial cell densities in the original Horonobe groundwater samples and incubated media were used to determine the extents to which cells were concentrated in the incubated media. The concentration factors (7.2 and 3.2 times for ESB214 and VSB249, respectively) were used to convert the AOM rates per 1 litter of the original groundwater. As a result, amorphous Fe(III) amendment with ESB214 showed relatively high AOM rates at 93.7 ± 40.6 nM/day. We also incubated ESB214 amended with nitrate and sulfate and without any oxidants.

**Fig. 3.**
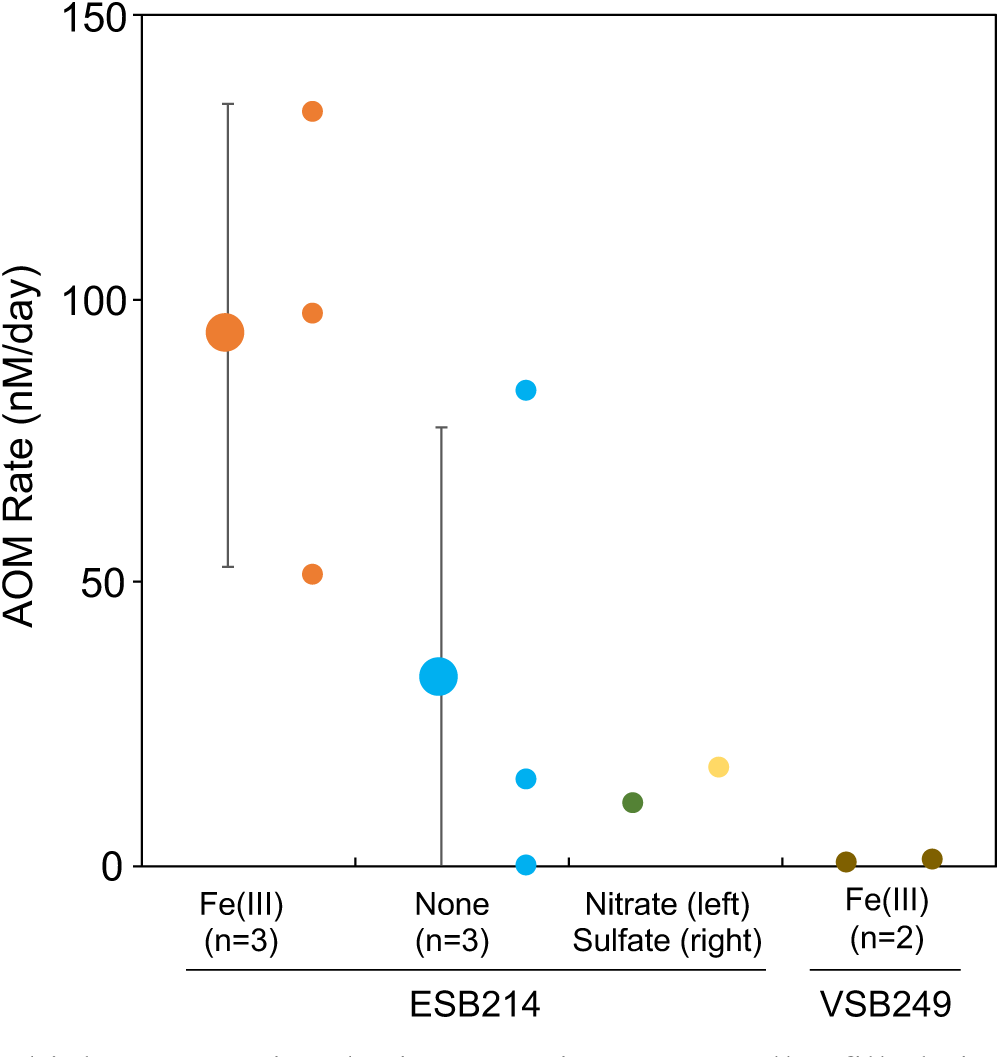
AOM rates obtained from high-pressure incubation experiments. Smaller filled circles show AOM rates for single experiments and larger ones with error bars indicate the means and standard deviations of the replication. The amended electron acceptors and replication numbers (*n*) are labelled with the circles and groundwater sample IDs.

These incubation experiments showed relatively low AOM rates, except for one data point obtained without the oxidant amendment (Fig. 3). With respect to VSB249, amorphous Fe(III) amendment was associated with nearly undetected levels of AOM activities.

### Characterization of suspended particulates in groundwater samples

These results suggest the presence of Fe(III) in ESB214, which is indicated by the high concentration of total iron relative to that of Fe(II) (Table 1). In addition, we visually recognized the loading of some yellowish-brown material on the filters after the passage of ESB214 (6.8 L). In contrast, the filters after the passage of VSB249 (64 L) gave no apparent loading on the filters. We analyzed the yellowish-brown material to evaluate the presence of insoluble Fe(III) for AOM. Sequential iron extraction method was first conducted as described in Yanase *et al*. (1991). After iron extraction, the ratio of Fe(III) relative to Fe(II) was determined by subtracting Fe(II) from total iron. The total iron content and the Fe(III) ratio were high in the amorphous iron (including ferrihydrite) and phyllosilicates (Fig. 4A), whereas those in carbonate and the crystalline iron were low. It is unlikely that the high reactivity of the amorphous iron exclude the possibility that the amorphous iron is originally present in ESB214 under reducing conditions (Eh = −219 mV) with dissolved Fe(II) (5 µM). Instead, amorphous Fe(III) was formed after the exposure of suspended particulates to air after sampling. Similarly, the high Fe(III) ratio in phyllosilicates was resulted artificially after sampling. Some smectite group phyllosilicates containing iron Fe(III) in their sheet structures are known to have the range of Fe(II)/Fe(III) ratio corresponding to surrounding redox potentials (Gorski *et al*., 2013). For instance, iron-bearing smectite minerals retain significant portions of Fe(III) even at ∼219 mV. Thus, the phyllosilicates in the suspended particles were further investigated mineralogically with X-ray diffractometry (XRD) and scanning electron microscopy equipped with energy dispersive spectrometry (SEM-EDS). The obtained EDS spectrum exhibited the elemental composition corresponding to a smectite mineral and a trace amount of iron in the structure (Fig. 4B). While the peak of smectite basal spacing and its shift after glycolation treatment reflecting the expansion of d_001_ were confirmed in XRD patterns (Fig. 4C). At the higher angle of the pattern, a peak of smectite (060) reflection suggested a dioctahedral sheet structure. To conclude, these results confirmed the presence of iron-bearing dioctahedral smectite in ESB214.

**Fig. 4.**
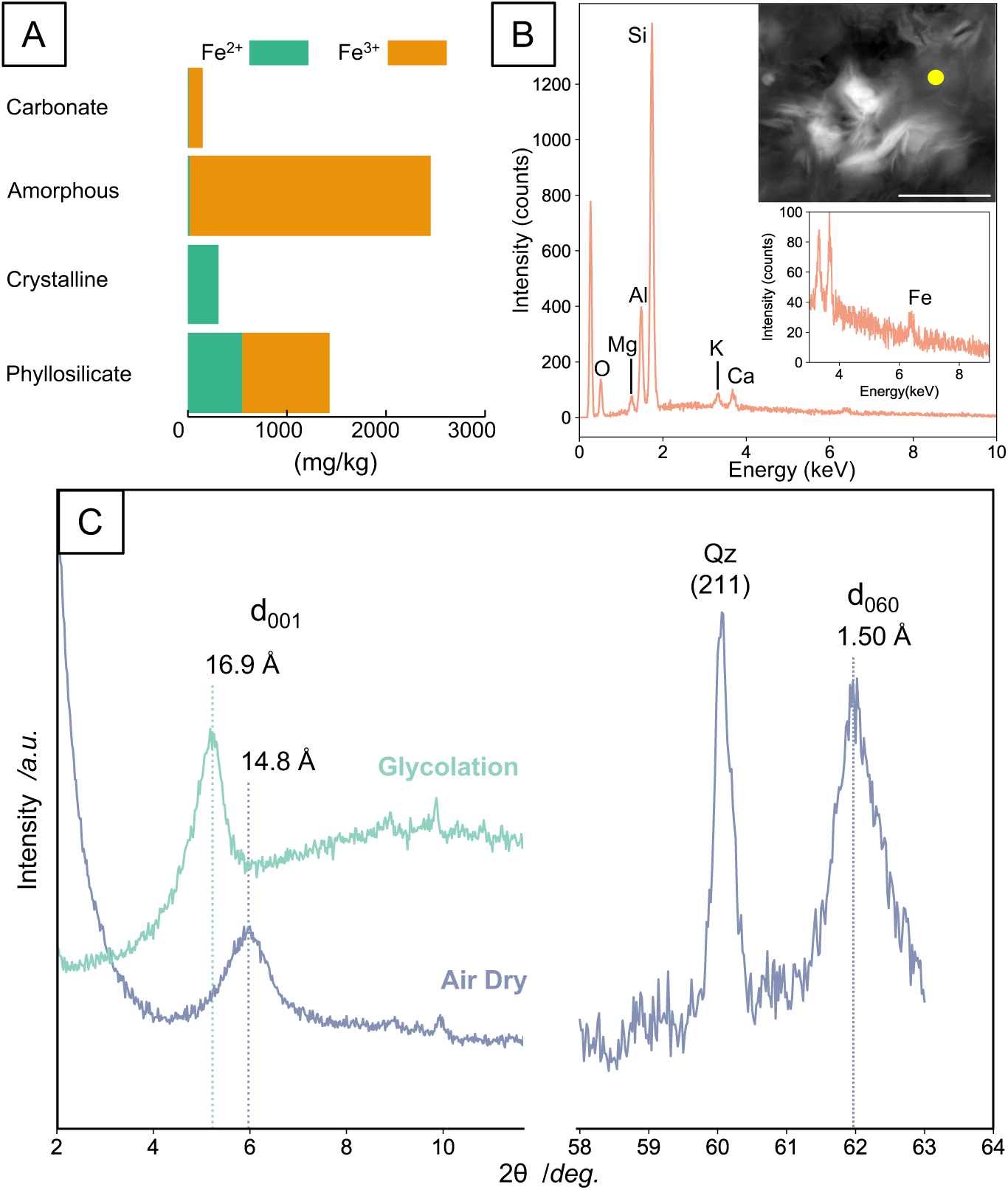
Mineralogical characteristics of suspended particulates in ESB214. Concentrations of ferric and ferrous iron in each extraction fraction of suspended particulates (A). Energy dispersive X-ray spectrum of the particulates and back-scattered electron image with yellow circle, where the spectrum was obtained (B). The scale bar in an electron microscopy image indicates 3.0 µm. Enlarged spectrum emphasizing a Fe peak is also shown. Powder X-ray diffraction (XRD) pattern of the particulates (C). Low-angle XRD patterns (2*θ*: 2-12°) of air-dried and ethylene glycol-sprayed samples for clarifying the expansion of basal spacings (left). High-angle pattern (2*θ*: 58-63°) for air-dried sample including (060) reflection of phyllosilicate minerals (right) for distinguishing dioctahedral and trioctahedral sheet structures.

## Discussion

### Subsurface AOM activities mediated by Ca. Methanoperedenaceae

The members of family *Ca*. Methanoperedenaceae have been reported from the deep terrestrial subsurface as well as to near-surface settings. Metabolic details of the near-surface members of *Ca*. Methanoperedenaceae have been characterized by incubation experiments under ambient pressure (Arshad *et al*., 2015; Vaksmaa *et al*., 2017; Weber *et al*., 2017; Guerrero-Cruz *et al*., 2018; Berger *et al*., 2021). However, metabolic characteristics of deep-subsurface members remain largely unknown.

In this study, we attempted to accurately measure AOM rates under in-situ high-pressure conditions using the newly developed laboratory incubation system. The incubation was set to mimic in-situ conditions as much as possible. A factor slightly different from the in-situ counterparts was the cell densities 7.2 and 3.2 times higher in the incubation media than in the original groundwater samples. Although the higher cell density might enhance the degradation of necromass, this effect was minimized by keeping the incubation time short. Nevertheless, there is the possibility that the AOM rates were underestimated, given that other Fe(III)-reducing activities (e.g. *Ferribacterium* spp.) are competitive for electron acceptors available in the incubation media. Even though the obtained rates might be slower, our data appear to be informative for the extent of the in-situ AOM activities and the electron acceptors in the borehole

From our incubation experiments, the highest AOM rates were obtained when amorphous Fe(III) was added to the incubation media prepared from ESB214. The Fe(III)-dependent AOM of *Ca*. Methanoperedenaceae archaea in ESB214 is hypothesized by our previous study, given that canonical correspondence analysis statistically clarified the relationship between the high concentration of total iron and the high abundance of *Ca*. Methanoperedenaceae (Hernsdorf *et al*., 2017). This relationship is also supported by results from *Ca*. Methanoperedenaceae-dominated bioreactor experiments which demonstrated ferrihydrite-dependent AOM activities (Ettwig *et al*., 2016; Cai *et al*., 2018). Therefore, the AOM activities observed in the incubation experiments serve as the first direct evidence that *Ca*. Methanoperedenaceae mediates AOM under subsurface conditions. This notion is also supported by the dominance of *Ca*. Methanoperedenaceae in the original ESB214 potentially as a sole anaerobic methanotroph.

### Potential in-situ electron acceptors for AOM

AOM activities measured by the incubation experiments are considered to reflect AOM activities in the boreholes. Nearly undetected AOM activities from the incubation experiments with VSB249 indicate substantially low AOM activities in the borehole. In contrast, moderately high AOM activities measured by the incubation experiments with ESB214 indicate that elevated levels of AOM activities are sustained in the borehole. Thus, it is critical to clarify electron acceptors such as Fe(III) available in the borehole for understanding factors controlling subsurface AOM activities.

Given the low stability of amorphous Fe(III) in ESB214, it is likely that the particulate component other than amorphous Fe(III) could be utilized as insoluble electron acceptors for AOM activities. Indeed, the particulates were found to contain Fe(III)-bearing smectite clay (Fig. 4A). Thus, detected levels of AOM activities without any amendment with the artificial oxidants were explained in ESB214 (“None” in Fig. 3). Similarly, AOM rates were also reasonable with the amendment of nitrate or sulfate (Fig. 3), given the availability of Fe(III)-bearing smectite clay in ESB214. Although the usage of nitrate and sulfate has been addressed for the respiration of the *Ca*. Methanoperedenaceae members (Haroon *et al*., 2013; Weber *et al*., 2017; Ino *et al*., 2018), these oxidants are scarce in Horonobe groundwater samples (Table 1). Hence, it is likely that equivalent AOM rates obtained from nitrate-or sulfate-amended incubation experiments were also dependent on particulates originally present in the borehole.

The importance of the particulates was indirectly evaluated for VSB249. The particulates were not visible on a 47-mm diameter filter after filtrating a large volume of the VSB249 groundwater. Our previous metagenomic analysis has shown that the *Ca*. Methanoperedenaceae members are nearly absent in VSB249 (Hernsdorf *et al*., 2017), which might be resulted from the lack of the particulates.

As expected, the AOM rate measurements of VSB249 revealed the slowest AOM rates even with the amendment of amorphous-Fe(III). This result highlights the importance of the particulates for the subsurface AOM activities.

It is still unknown whether *Ca*. Methanoperedenaceae archaea can utilize Fe(III) in phyllosilicate minerals. Previously, the capability of Fe(III) reduction in smectite minerals has been demonstrated for the bacterial groups with high numbers of MHCs gene in their genomes such as the genera *Shewanella* and *Geobacter* (Dong *et al*., 2009, and references therein). Thus, the possibility of electron transfer to Fe(III) in the smectite clay found in the particulates by the members of *Ca*. Methanoperedenaceae with the high number of MHCs gene will be investigated.

### Implications for geological disposal of nuclear wastes

AOM activities coupled to Fe(III) reduction results in abiotic reduction of U(VI) by biologically reduced iron (Wu *et al*., 2010). Likewise, ^99^Tc, a long-lived fission product of ^235^U, is reduced by Fe(II) (De Luca *et al*., 2001). Enzymatic reduction of U(VI) is expected, given the abundance of MHCs in the genomes (Marshall *et al*., 2006). Thus, the subsurface migration of long-lived radionuclides is profoundly influenced by AOM activities demonstrated in the borehole. Stimulation of AOM activities coupled to Fe(III) reduction after back-filling of deep repositories for high-level nuclear wastes leads to the establishment of reducing conditions and the retardation of radionuclide migration.

## Conclusion

In this study, *Ca*. Methanoperedenaceae-abundant groundwater was subjected to in-situ pressure incubation with ^13^C-labelled methane. Without the amendments of electron acceptors, AOM activity was experimentally demonstrated. In addition, Fe(III) available from a smectite clay mineral in suspended particles to might serve as electron acceptors in the borehole. Our data conclusively show the mediation of AOM in the deep subsurface borehole.

## Acknowledgements

The Ministry of Economy, Trade and Industry of Japan has funded a part of the work as “The project for validating near-field system assessment methodology in geological disposal system” (2020 FY, Grant Number: JPJ007597).We acknowledge technical supports from Koji Ichimura.

## Materials and Methods

### Site description and biological sampling

The Horonobe URL was constructed by Japan Atomic Energy Agency with the objectives of research and development of geological disposal technologies of high-level radioactive waste. The sediments around the URL consist of the Wakkanai Formation (Neogene siliceous mudstones) overlain by the Koetoi Formation (Neogene to Quaternary diatomaceous mudstones). The origin of ESB214 and VSB249 are fossil seawater mostly diluted by meteoric water (Teramoto *et al*., 2006; Iwatsuki *et al*., 2009). Groundwater samples were filtered with a membrane filter (type GPWP, 0.22 µm pore size: Merck Millipore, Darmstadt, Germany) in a pressure-resistant stainless filter holder (HP Filter Holder, 47 mm, stainless steel: Merck Millipore, Darmstadt, Germany). Filtered groundwater samples were obtained from the filter holder directly into Ar-flushed 100-ml glass vials. The filter holders and the filtered groundwater were stored at 4 °C.

### High-pressure incubation

The amorphous Fe(III) was synthesized by titrating 80 mM FeCl_3_·6H_2_O solution by 10 N NaOH and thoroughly rinsed with deionized water before amendment. ^13^C-labeled methane (CLM-3590; Cambridge Isotope Laboratories, Andover, MA, USA) was stored inside Tedlar bags for one week before amendment to remove trace amount of H_2_ contained as impurities.

The media with cell suspensions were prepared as described above. The media were originally prepared in 100 mL glass vials sealed with a butyl rubber stopper and an aluminum cap. After preparation other than methane amendment, Ar-gas purging for each vial was conducted repeatedly until the oxygen level became lower than the detection limit of fiber-optic oxygen meter (MICROX TX3-TRACE; PreSens, Regensburg, Germany). The media were transferred into Ar-purged Tedlar bags inside a glove box, and the treated ^13^C-labeled methane was amended for each bag with a gas-tight syringe. The bags were transferred into a stainless vessel. The original gas phase in the stainless vessel was replaced and pressurized with Ar gas.

Some portions of the media with cell suspensions were fixed with 3.7% formalin solution at neutral pH for cell counting and filtered with 0.2-µm pore-sized membrane filters. Filtrates were stored at 4 °C in blood-collection tubes sealed with thick butyl rubber stoppers to prevent air exchange and served for *δ*^13^C analysis. The remaining media were stored in plastic tubes for other chemical analyses at 4 °C. The media with cell suspensions were also processed at the beginning of the incubation.

### Chemical analysis procedures

DIC concentrations were estimated by measuring alkalinity in the filtrates by acid titration using 0.02-N hydrochloric acid. Based on the amount of 0.02-N hydrochloric acid and pH measured by a pH meter (LAQUA F-73, Horiba, Japan), alkalinity was calculated by a Gran plot method.

The filtrated samples stored inside of blood-collection tubes were transferred into glass vials sealed with rubber stopper and aluminum cap containing sulfamic acid. After the reaction with the acid, DIC species, mainly bicarbonate, were extracted as carbon dioxide. The *δ*^13^C of carbon dioxide were analyzed with an IsoPrime100 isotope ratio mass spectrometer (Isoprime Ltd., Cheadle Hulme, UK) equipped with a customized continuous-flow gas preparation system (MICAL3c; Ishimura *et al*., 2004;2008) at Kyoto University, Japan.

### Microscopic observations

We conducted direct cell counting for measurement of total cell numbers in each incubated medium and original groundwater sample. In detail, 0.1 mL of fixed cell suspension was filtrated by 0.22-µm pore-sized black polycarbonate filter (type GTBP; Merck Millipore, Darmstadt, Germany) and stained with 10 × SYBR Green I (SYBR^®^ Green I Nucleic Acid Stain; Lonza, Rockland, ME, USA) in 1×TAE buffer for 3 min. at room temperature. Microbial cells on the filter were directly counted using an epifluorescence microscope (BX51; Olympus, Tokyo, Japan) equipped with a digital camera (DP70; Olympus, Tokyo, Japan). Two filters per sample were prepared and cell numbers were obtained by observation of 50 fields of view on a single specimen and calculation of their average.

### DNA extraction, sequencing and phylogenetic analysis

ESB214 was filtered on a 0.22-μm pore size filter (type GVWP; Millipore) and subjected to DNA extraction using MO BIO’S PowerMax Soil DNA Isolation Kit (Qiagen, Inc., Valencia, CA, USA). 16S rRNA gene sequences were amplified by polymerase chain reaction (PCR) using LA *Taq* polymerase (TaKaRa-Bio, Inc., Kusatsu, Japan) for Illumina MiSeq paired-end sequencing. The primers Uni530F and Uni907R (Nunoura *et al*., 2012) containing Illumina TruSeq adapter sequences (Illumina Inc., San Diego, CA, USA) were used for PCR. A reaction mixture was prepared in which the concentration of each oligonucleotide primer was 0.1 μM and that of the DNA template was ca. 0.1 ng/μL. Thermal cycling was performed with 35 cycles of denaturation at 96 °C for 20 s, annealing at 56 °C for 45 s, and extension at 72 °C for 120 s. The first PCR amplicon was used for the second PCR step, which was run with TruSeq P5 and Index-containing P7 adapters, and PCR program identical to the first, except with 10 cycles. The final PCR amplicon with the expected size was confirmed by electrophoresis on TAE (40 mM Tris acetate, 1 mM EDTA, pH 8.3) agarose gels (1 %), which was purified using a MinElute Gel Extraction Kit (Qiagen, Inc., Valencia, CA, USA). 16S rRNA gene sequencing was performed using a MiSeq platform with MiSeq Reagent Kit v2 (Illumina).

The paired-end sequence reads were demultiplexed, trimmed and quality filtered. After the removal of chimeras within BaseSpace (Illumina). The screened reads were clustered with a 97% sequence similarity cut-off using Mothur (Schloss *et al*., 2009), and the output consensus (Operational Taxonomic Units; OTUs) sequences were used as phylotypes. For alignment and taxonomic affiliation of OTUs, SILVA SSU Ref NR database was used (Quast et al., 2013). The neighbor-joining tree based on 16S rRNA gene sequences were constructed in the ARB software (Ludwig *et al*., 2004) for *Ca*. Methanoperedenaceae-affiliated OTUs and closely related sequences retrieved from GenBank (http://www.ncbi.nlm.nih.gov/genbank/). Distantly related sequences were retrieved from The SILVA SSU Ref NR database for the tree construction. Bootstrap analysis of the maximum-likelihood tree was performed with 1000 replicates in the Geneious software package (Kearse *et al*., 2012).

### Geochemical and mineralogical analyses for suspended particulates

Sequential extraction of iron from suspended particulates in ESB214 was conducted for quantification of ferric and ferrous iron. In details, the suspended particulates were treated with (1) 1 M sodium acetate (pH 5.0) for 4 hours and (2) 12.6 g/L of oxalic acid and 18.4 g/L of ammonium oxalate for 4 hours at room temperature. The iron in carbonate and adsorbed phases from (1), amorphous or poorly crystalline phases from (2) were extracted separately. The residue was further treated with (3) 17.4 g/L sodium dithionite, 0.3 M trisodium citrate and 0.2 M sodium hydrogen carbonate for 60 min. and (4) 6 M hydrochloric acid for 150 min. at 85 °C. These steps extracted iron from crystalline minerals (3) and from clay minerals (4), respectively. After each treatment, the samples were centrifuged at 3500 rpm for 30 min. The precipitate was rinsed with deionized water before next treatment, while the supernatant was served for concentration measurements. The ferrous iron in supernatant was reacted with phenanthroline, and the concentration was calculated from absorbance at 510 nm measured by a spectrophotometer (U2900; Hitachi, Tokyo, Japan). Afterwards, the specimen was amended with hydrochloric acid, and the total iron concentration was measured by an atomic absorption spectrophotometer (ZA3300; Hitachi, Tokyo, Japan). The ferric iron concentration was obtained by subtraction of ferrous iron concentration from total iron. These spectrometric procedures were repeated at least three times for each sample.

The phyllosilicate mineral characterization was performed by XRD pattern analysis. The suspended clay-sized particulates from ESB214 were obtained and vacuum-dried. The clay-sized particulates were mounted on a non-reflecting sample holder. The diffraction pattern was obtained with monochromatized Cu-Kα X-ray at an operation voltage of 40 kV and an operation current of 30 mA using an X-ray diffractometer (RINT-2100; Rigaku, Japan). After ethylene glycol was sprayed, diffraction pattern of the wet sample was also obtained in addition to the dried sample. The 2*θ* range and resolution were 5 – 65° and 0.02°, respectively.

The portion of the dried clay-sized particulates were embedded in LR White Resin (London Resin Co., Ltd., Aldermaston, England) and solidified in an oven at 50 °C for 48h. The resin block was coated with carbon (SC-701C Quick Carbon Coater; Sanyu Denshi Co., Ltd., Tokyo, Japan), and the morphological observation of suspended particulates and their elemental composition were obtained by a field emission SEM (S-4500; Hitachi, Tokyo, Japan) equipped with an EDS detector (Ultradry EDS detector NS7; Thermo Fischer Scientific Inc., Waltham, MA, USA). The analysis was operated at accelerating voltage of 15 kV and emission current of 13 µA.

## References

Arshad, A., Speth, D.R., de Graaf, R.M., Op den Camp, H.J., Jetten, M.S., and Welte, C.U. (2015) A metagenomics-based metabolic model of nitrate-dependent anaerobic oxidation of methane by Methanoperedens-like archaea. Front Microbiol 6: 1423.

Berger, S., Cabrera-Orefice, A., Jetten, M.S., Brandt, U., and Welte, C.U. (2021) Investigation of central energy metabolism-related protein complexes of ANME-2d methanotrophic archaea by complexome profiling. BBA-Bioenergetics 1862: 148308.

Boetius, A., Ravenschlag, K., Schubert, C.J., Rickert, D., Widdel, F., Gieseke, A. et al. (2000) A marine microbial consortium apparently mediating anaerobic oxidation of methane. Nature 407: 623–626.

Cai, C., Leu, A.O., Xie, G.-J., Guo, J., Feng, Y., Zhao, J.-X. et al. (2018) A methanotrophic archaeon couples anaerobic oxidation of methane to Fe (III) reduction. ISME J 12: 1929–1939.

Cummings, D.E., Caccavo Jr, F., Spring, S., and Rosenzweig, R.F. (1999) Ferribacterium limneticum, gen. nov., sp. nov., an Fe (III)-reducing microorganism isolated from mining-impacted freshwater lake sediments. Arch Microbiol 171: 183–188.

Dong, H., Jaisi, D.P., Kim, J., and Zhang, G. (2009) Microbe-clay mineral interactions. Am Mineral 94: 1505–1519.

Ettwig, K.F., Zhu, B., Speth, D., Keltjens, J.T., Jetten, M.S., and Kartal, B. (2016) Archaea catalyze iron-dependent anaerobic oxidation of methane. P Natl Acad Sci USA 113: 12792–12796.

Ettwig, K.F., Butler, M.K., Le Paslier, D., Pelletier, E., Mangenot, S., Kuypers, M.M. et al. (2010) Nitrite-driven anaerobic methane oxidation by oxygenic bacteria. Nature 464: 543–548.

Flynn, T.M., Sanford, R.A., Ryu, H., Bethke, C.M., Levine, A.D., Ashbolt, N.J., and Santo Domingo, J.W. (2013) Functional microbial diversity explains groundwater chemistry in a pristine aquifer. BMC Microbiol 13: 1–15.

Gorski, C.A., Klüpfel, L.E., Voegelin, A., Sander, M., and Hofstetter, T.B. (2013) Redox properties of structural Fe in clay minerals: 3. Relationships between smectite redox and structural properties. Environ Sci Technol 47: 13477–13485

Guerrero-Cruz, S., Cremers, G., van Alen, T.A., Op den Camp, H.J., Jetten, M.S., Rasigraf, O., and Vaksmaa, A. (2018) Response of the anaerobic methanotroph “Candidatus Methanoperedens nitroreducens” to oxygen stress. Appl Environ Microb 84: e01832–01818.

Haroon, M.F., Hu, S., Shi, Y., Imelfort, M., Keller, J., Hugenholtz, P. et al. (2013) Anaerobic oxidation of methane coupled to nitrate reduction in a novel archaeal lineage. Nature 500: 567–570.

Hernsdorf, A.W., Amano, Y., Miyakawa, K., Ise, K., Suzuki, Y., Anantharaman, K. et al. (2017) Potential for microbial H<sub>2</sub> and metal transformations associated with novel bacteria and archaea in deep terrestrial subsurface sediments. ISME J 11: 1915–1929.

Hinrichs, K.-U., Hayes, J.M., Sylva, S.P., Brewer, P.G., and DeLong, E.F. (1999) Methane-consuming archaebacteria in marine sediments. Nature 398: 802–805.

Ino, K., Hernsdorf, A.W., Konno, U., Kouduka, M., Yanagawa, K., Kato, S. et al. (2018) Ecological and genomic profiling of anaerobic methane-oxidizing archaea in a deep granitic environment. ISME J 12: 31–47.

Ishimura, T., Tsunogai, U., and Gamo, T. (2004) Stable carbon and oxygen isotopic determination of sub-microgram quantities of CaCO<sub>3</sub> to analyze individual foraminiferal shells. Rapid Commun Mass Sp 18: 2883–2888.

Ishimura, T., Tsunogai, U., and Nakagawa, F. (2008) Grain-scale heterogeneities in the stable carbon and oxygen isotopic compositions of the international standard calcite materials (NBS 19, NBS 18, IAEA-CO-1, and IAEA-CO-8). Rapid Commun Mass Sp 22: 1925–1932.

Iwatsuki, T., Ishii, E., and Niizato, T. (2009) Scenario development of long-term evolution for deep hydrochemical conditions in Horonobe area, Hokkaido, Japan. J Geogr 118: 700–716.

Kearse, M., Moir, R., Wilson, A., Stones-Havas, S., Cheung, M., Sturrock, S. et al. (2012) Geneious Basic: an integrated and extendable desktop software platform for the organization and analysis of sequence data. Bioinformatics 28: 1647–1649.

Kletzin, A., Heimerl, T., Flechsler, J., van Niftrik, L., Rachel, R., and Klingl, A. (2015) Cytochromes c in Archaea: distribution, maturation, cell architecture, and the special case of Ignicoccus hospitalis. Front Microbiol 6: 439.

Krüger, M., Wolters, H., Gehre, M., Joye, S.B., and Richnow, H.-H. (2008) Tracing the slow growth of anaerobic methane-oxidizing communities by <sup>15</sup>N-labelling techniques. FEMS Microbiol Ecol 63: 401–411.

Leu, A.O., McIlroy, S.J., Ye, J., Parks, D.H., Orphan, V.J., and Tyson, G.W. (2020a) Lateral gene transfer drives metabolic flexibility in the anaerobic methane-oxidizing archaeal family Methanoperedenaceae. MBio 11: e01325–01320.

Leu, A.O., Cai, C., McIlroy, S.J., Southam, G., Orphan, V.J., Yuan, Z. et al. (2020b) Anaerobic methane oxidation coupled to manganese reduction by members of the Methanoperedenaceae. ISME J 14: 1030–1041.

Lu, Y.-Z., Fu, L., Ding, J., Ding, Z.-W., Li, N., and Zeng, R.J. (2016) Cr (VI) reduction coupled with anaerobic oxidation of methane in a laboratory reactor. Water Res 102: 445–452.

Ludwig, W., Strunk, O., Westram, R., Richter, L., Meier, H., Yadhukumar et al. (2004) ARB: a software environment for sequence data. Nucleic Acids Res 32: 1363–1371.

De Luca, G., de Philip, P., Dermoun, Z., Rousset, M., and Verméglio, A. (2001) Reduction of technetium (VII) by Desulfovibrio fructosovorans is mediated by the nickel-iron hydrogenase. Appl Environ Microb 67: 4583–4587.

Mardanov, A.V., Kadnikov, V.V., Beletsky, A.V., and Ravin, N.V. (2020) Sulfur and methane-oxidizing microbial community in a terrestrial mud volcano revealed by metagenomics. Microorganisms 8: 1333.

Methé, B., Nelson, K.E., Eisen, J.A., Paulsen, I.T., Nelson, W., Heidelberg, J. et al. (2003) Genome of Geobacter sulfurreducens: metal reduction in subsurface environments. Science 302: 1967–1969.

Marshall, M.J., Beliaev, A.S., Dohnalkova, A.C., Kennedy, D.W., Shi, L., Wang, Z. et al. (2006) c-Type cytochrome-dependent formation of U (IV) nanoparticles by Shewanella oneidensis. PLoS Biol 4: e268.

Mezawa, T., Mochizuki, A., Miyakawa, K., and Sasamoto, H. (2018) Records of physico-chemical parameters by geochemical monitoring system in the Horonobe Underground Research Laboratory. Available at https://doi.org/10.11484/jaea-data-code-2018-001

Miettinen, H., Bomberg, M., and Vikman, M. (2018) Acetate activates deep subsurface fracture fluid microbial communities in Olkiluoto, Finland. Geosciences 8: 399.

Miyakawa, K., Ishii, E., Hirota, A., Komatsu, D.D., Ikeya, K., and Tsunogai, U. (2017) The role of low-temperature organic matter diagenesis in carbonate precipitation within a marine deposit. Appl Geochem 76: 218–231.

Miyakawa, K., Mezawa, T., Mochizuki, A., and Sasamoto, H. (2020) Data of groundwater chemistry obtained in the Horonobe Underground Research Laboratory Project (FY2017-FY2019). Available at https://doi.org/10.11484/jaea-data-code-2020-001

Nauhaus, K., Boetius, A., Krüger, M., and Widdel, F. (2002) In vitro demonstration of anaerobic oxidation of methane coupled to sulphate reduction in sediment from a marine gas hydrate area. Environ Microbiol 4: 296–305.

Nanjo, I., Amano, Y., Iwatsuki T., Kunimaru, T., Murakami, H., Hosoya, S., and Morikawa, K. (2012) Development of a groundwater monitoring system at Horonobe Underground Research Center. Available at https://doi.org/10.11484/jaea-research-2011-048

Nunoura, T., Takaki, Y., Kazama, H., Hirai, M., Ashi, J., Imachi, H., and Takai, K. (2012) Microbial diversity in deep-sea methane seep sediments presented by SSU rRNA gene tag sequencing. Microbes Environ 27: 382–390.

Quast, C., Pruesse, E., Yilmaz, P., Gerken, J., Schweer, T., Yarza, P. et al. (2013) The SILVA ribosomal RNA gene database project: improved data processing and web-based tools. Nucleic Acids Res 41: D590–D596.

Schloss, P.D., Westcott, S.L., Ryabin, T., Hall, J.R., Hartmann, M., Hollister, E.B. et al. (2009) Introducing mothur: open-source, platform-independent, community-supported software for describing and comparing microbial communities. Appl Environ Microb 75: 7537–7541.

Shi, L., Squier, T.C., Zachara, J.M., and Fredrickson, J.K. (2007) Respiration of metal (hydr) oxides by Shewanella and Geobacter: a key role for multihaem c-type cytochromes. Mol Microbiol 65: 12–20.

Shi, L., Dong, H., Reguera, G., Beyenal, H., Lu, A., Liu, J. et al. (2016) Extracellular electron transfer mechanisms between microorganisms and minerals. Nat Rev Microbiol 14: 651–662.

Teramoto, M., Shimada, J., and Kunimaru, T. (2006) Evidences of groundwater regime in impermeable rocks by stable isotopes in porewaters of drilled cores. J Jpn Soc Eng Geol 47: 68–76.

Timmers, P.H., Gieteling, J., Widjaja-Greefkes, H.A., Plugge, C.M., Stams, A.J., Lens, P.N., and Meulepas, R.J. (2015) Growth of anaerobic methane-oxidizing archaea and sulfate-reducing bacteria in a high-pressure membrane capsule bioreactor. Appl Environ Microb 81: 1286–1296.

Vaksmaa, A., Guerrero-Cruz, S., van Alen, T.A., Cremers, G., Ettwig, K.F., Lüke, C., and Jetten, M.S. (2017) Enrichment of anaerobic nitrate-dependent methanotrophic ‘Candidatus Methanoperedens nitroreducens’ archaea from an Italian paddy field soil. Appl Microbiol Biot 101: 7075–7084.

Weber, H.S., Habicht, K.S., and Thamdrup, B. (2017) Anaerobic methanotrophic archaea of the ANME-2d cluster are active in a low-sulfate, iron-rich freshwater sediment. Front Microbiol 8: 619.

Wu, W.-M., Carley, J., Green, S.J., Luo, J., Kelly, S.D., Nostrand, J.V. et al. (2010) Effects of nitrate on the stability of uranium in a bioreduced region of the subsurface. Environ Sci Technol 44: 5104–5111.

Yanase, N., Nightingale, T., Payne, T., and Duerden, P. (1991) Uranium distribution in mineral phases of rock by sequential extraction procedure. Radiochim Acta 52-53: 387–394.

